# Mobilization of a *Kluyvera*-like *arnBCADTEF* operon, besides *mcr* genes, confers colistin resistance to *Escherichia coli* isolated from healthy animals

**DOI:** 10.1101/2020.08.06.240812

**Authors:** Alejandro Gallardo, María-Rocío Iglesias, María Ugarte-Ruiz, Marta Hernández, Pedro Miguela-Villoldo, Gloria Gutiérrez, David Rodríguez-Lázaro, Lucas Domínguez, Alberto Quesada

## Abstract

The use of colistin as a last resort antimicrobial is compromised by the emergence of resistant enterobacteria with acquired determinants like *mcr* genes, mutations that activate the PmrAB two-component system and also by some other(s) still unknown mechanism(s). This work analyzed 74 *E. coli* isolates from healthy swine, turkey or bovine animals, characterizing their colistin resistance determinants. The *mcr-1* gene, detected in 69 isolates, was the main determinant found among which 45% were carried by highly mobile plasmids, followed by four strains lacking previously known resistance determinants or two with *mcr-4* (one in addition to *mcr-1*), whose phenotypes were not transferred by conjugation. Although a fraction of isolates carrying *mcr-1* or *mcr-4* genes also presented missense polymorphisms in *pmrA* or *pmrB*, constitutive activation of PmrAB was not detected, in contrast to control strains carrying mutations that confer colistin resistance. The expression of *mcr* genes negatively controls *arnBCADTEF* expression, a down-regulation that was also observed in the four isolates lacking known resistance determinants, three of them sharing the same macrorestriction and plasmid profiles. Genomic sequencing of one of these strains, isolated from a bovine in 2015, revealed a IncFII plasmid of 60 Kb encoding an *arnBCADTEF* operon closely related to *Kluyvera ascorbata* homologs. This element, named pArnT1, was cured by ethidum bromide and lost in parallel to colistin resistance. This work reveals that, besides *mcr* genes and chromosomal mutations, mobilization of *arnBCADTEF* operon represents a colistin resistance mechanism whose spread and relevance for public health should be carefully surveyed.

**Abstract Importance:** Colistin is an old antibiotic that has returned to first-line fighting against (Gram negative) microorganisms after pandemic rising of antimicrobial resistance. However, low susceptibility to colistin is also becoming spread, mainly by plasmid mobilization of one of the enzymes (encoded by *mcr* genes) that modify covalently the external layer (the lipid A component of the lipopolysaccharide) of bacterial envelope, interfering antibiotic effectiveness. The second enzymatic system that performs envelope modification and confers colistin resistance when overexpressed is encoded by *arnBCADTEF* operon, a set of seven genes with location restricted (up to now) to the chromosome of Gram negative bacteria. This work describes plasmid mobilization of this operon between enterobacteria, from *Kluyvera* to *Escherichia coli*, where a *Kluyvera*-like *arnBCADTEF* operon carried by pArnT1 might represent, besides *mcr* genes, a potential risk for antimicrobial therapy and might require careful surveillance.

## Introduction

For a long time it was believed that Gram-negative bacteria could only acquire resistance to polymyxins B or E (colistin), two closely-related cationic and cyclic antimicrobial peptides, by mutations altering signaling processes that control covalent modifications of LPS and reduce its negative charge (Zhang et al., 2019). Polymorphisms that render constitutively active PmrAB or PhoPQ two-component systems (TCS) lead to overexpression of two enzymes, EptA (PmrC) and ArnT (PmrK), that transfer phosphoethanoamine (PetN) or 4-amino-4-deoxy-L-arabinose (L-Ara4N), respectively, to one or both phosphates bounded to disaccharide of lipid A and reduce colistin binding (Figure 1). Encoded by operons *eptApmrAB* and *arnBCADTEF* (*pmrHFIJKLM*), the deregulated expression from these genes interferes with cell viability, making colistin resistant mutants low prevalent under non selective conditions and reducing the risk of antimicrobial resistance spread (Olaitan et al., 2014). Therefore, until very recently colistin was routinely used to control intestinal enterobacteria in farm animals, while confidence in its efficacy as the last resort antibiotic for control of insidious infections in human was maintained since vertical transmission of resistant clones was considered highly improbable and plasmidic determinants were yet unknown. However, during the last five years colistin resistance has been widely spread among enterobacteria by plasmids carrying different genes belonging to the *mcr*/*eptA* family. *mcr* genes originated by plasmid mobilization of *eptA* orthologs from different species, targeting the encoded transferase activity to the bacterial inner membrane where its phosphatidylethanolamine substrate is an abundant component of the phospholipid pool (Zhang et al., 2019). Together with their efficient transfer by a large number of promiscuous replicons, plasmids carrying *mcr* genes among which *mcr-1* is the most prevalent have managed to spread rapidly and widely among enterobacteriaceae, challenging the clinical utility of polymyxins.

**Figure 1.**
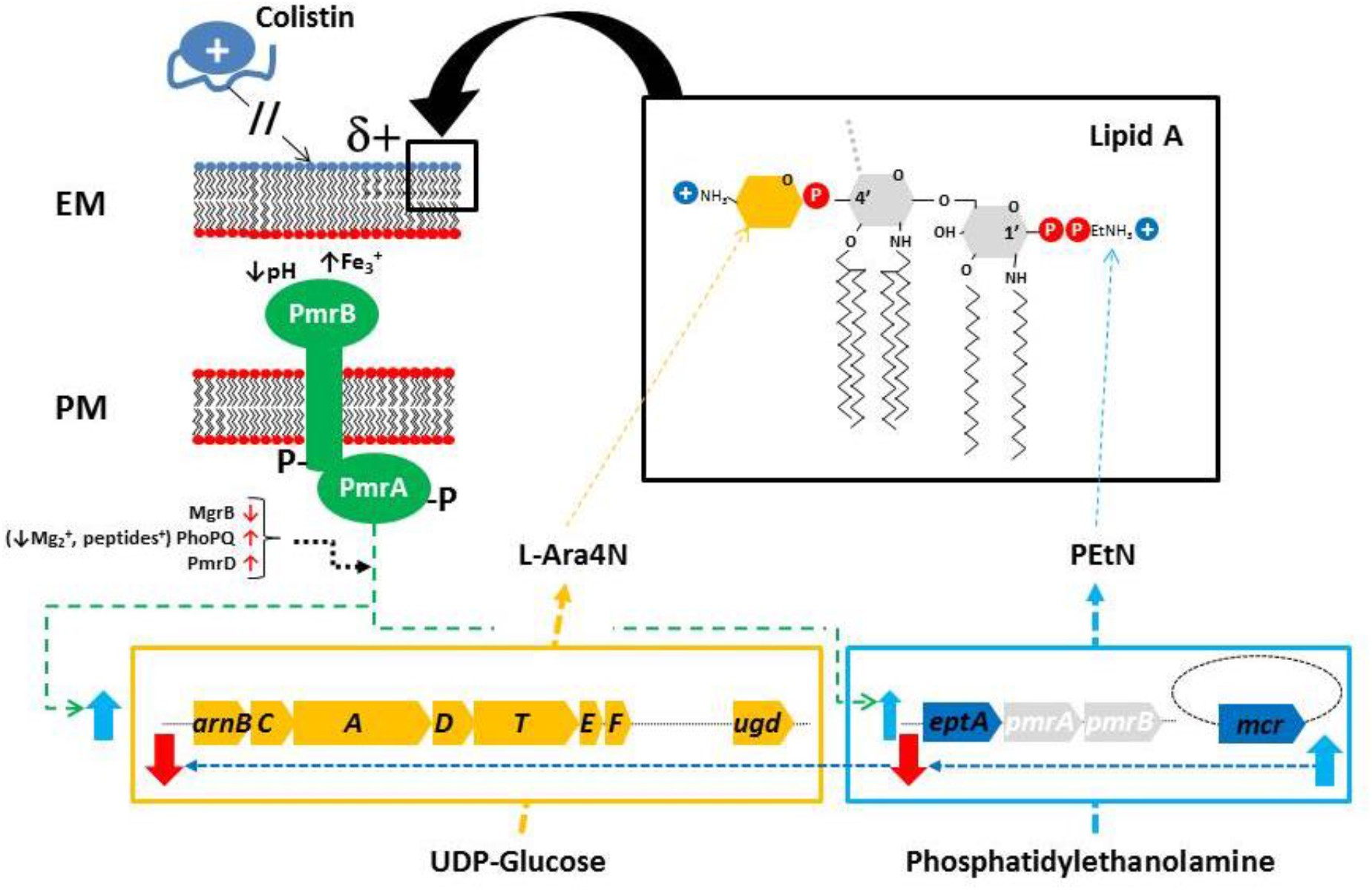
Expression of colistin resistance determinants in *E. coli.* The lipid A fraction of LPS and its major modifications by L-Ara4N and PEtN that block binding of colistin to the outer leaflet of external membrane (EM) are shown. The TCS PmrAB transduces low pH and high Fe^3+^ signals through the plasma membrane (PM) to the cytosol, where it activates transcription of effector genes. The possible role of the TCS PhoPQ mediating the activation of PmrAB in *E. coli* is controversial and might depend of particular genetic backgrounds (Olaitan et al., 2014), whereas expression of *mcr* genes down-regulates *arnBCADTEF* and *eptApmrAB* operons (this work).

In parallel to EptA/Mcr functions, the *arnBCADTEF* operon encodes the enzymes that synthesizes L-Ara4N from UDP-glucuronic acid and its transfer to 4’-phosphate lipid A (Figure 1), involving a seven-step biochemical pathway much more complex than the PEtN addition to lipid A by a single enzyme, probably explaining why the last is the only known plasmid determinant conferring colistin resistance (Zhang et al., 2019). The synthesis of L-Ara4N starts with multifunctional ArnA that catalyzes NAD^+^-dependent oxidative decarboxylation of UDP-glucuronic acid and formylation of UDP-L-Ara4N, whereas transamination of the ketopentose intermediate, addition of formylated L-Ara4N to undecaprenyl-P (Up) in the inner leaflet of the plasma membrane, deformylation of Up-L-Ara4FN, flip-flop of Up-L-Ara4N to the outer surface of the inner membrane and transfer of L-Ara4N from Up to lipid A, are performed by ArnB, ArnC, ArnD, ArnE/ArnF and ArnT, respectively (Raetz et al., 2007). Besides this, the Ugd (PmrE) enzyme is a UDP-glucose dehydrogenase that channels UDP-glucuronic acid for LPS modification, and it is encoded independently from *arnBCADTEF* genes although closely co-regulated by *pmrAB* (Olaitan et al., 2014).

In previous works we analyzed 7 colistin resistant *E. coli* isolates from healthy turkey, swine and bovine, finding out *mcr-1* in all strains, among which one carried an additional *mcr-3* gene in the same plasmid and two presented mutations conferring colistin resistance in *pmrA* or *pmrB* (Quesada et al., 2015; 2016; Hernández et al., 2017). These *mcr-1* genes were detected in plasmids with different sizes and, among them, 42.9% transferred efficiently colistin resistance by conjugation. In the present work we extend our screening to 74 additional strains, characterizing their colistin resistance determinants, among which *mcr-1* is by far the most prevalent. This larger strain collection contains strains lacking previously known colistin resistance determinants, some of which presented a plasmid carrying a *Kluyvera*-like *arnBCADTEF* operon that is associated with resistance to the antibiotic, a phenomenon evoking the spread of *mcr* genes and that could represent a second wave of plasmid-mediated colistin resistance elements which relevance should be carefully survey in next future.

## Materials and Methods

### Strains and growth conditions

In the context of the Spanish Surveillance Network of Antimicrobial Resistance in Bacteria of Veterinary Origin (VAV Network, VISAVET), *E. coli* strains isolated from turkey feces in 2014 and 2016, and from swine or bovine feces in 2015, have been included in the current study (supplementary Table I). Samples were pooled in situ from ten different animals in the case of turkey and from two different animals in the case of swine and bovine. All the samples were collected at slaughterhouse in the frame of samplings carried out in Spain, being randomly taken from samplings covering all the country and performed according to the rules of the EU Authorities (European Commission implementing decision 2013/652/EU).

### Determination of colistin resistance

Minimal inhibitory concentration (MIC) of colistin (mg/L) was determined by the 2-fold broth microdilution reference method according to ISO 20776-1:2006. The threshold of antimicrobial resistance adopted in this work was based on the epidemiological cut-off (ECOFF) values recommended by EUCAST (http://www.eucast.org). *E. coli* strain ATCC25922 was used as quality control of the technique.

### Screening colistin resistance determinants

Among the 10 plasmid *m*ediated *colistin resistance* determinants (*mcr* genes) (Wang et al., 2020), the first four genes *mcr-1* to *mcr-4* were screened by PCR with primers and conditions previously described (supplementary Table II and references therein). In addition, further screening was performed with a set of primers designed from sequence alignment of *mcr-1* to *mcr-8* genes (Zhang et al., 2019 and references therein) to specifically detect *mcr-1/-2/-6*, *mcr-5*, *mcr-3/-7* or *mcr-4/-8* in separated PCRs, whose functionality was evidenced by using DNA from control strains carrying *mcr-1/-2/-3/-4* or *mcr-5*. Recently described *mcr-9* and *mcr-10* genes have been voluntarily omitted from this PCR screening since, in contrast to all previously known *mcr* genes, its role in colistin resistance has yet to be demonstrated and could be marginal at best (Khedher et al., 2020; Wang et al., 2020).

The sequences from pmrAB genes were screened by PCR (supplementary Table II) in search of missense polymorphisms of encoded proteins, excluding those variants found in colistin susceptible strains, like *E. coli* ATCC25922, K12 or recently isolated from farm animals (Quesada et al., 2015). Primers were synthetized by STABVIDA (Caparica, Portugal), and sequencing of DNA fragments was performed in the facilities of the University of Extremadura, in the “Servicio de Técnicas Aplicadas a la Biociencia (STAB)”.

### Mobilization potential of colistin resistance determinants

Conjugations were performed by mating donor cells with *E. coli* J53 strain and selecting conjugants in 200 mg/L sodium azide and 2 mg/L colistin, and the efficiency was calculated dividing the number of conjugants by that of donor cells, sharing resistance to both compounds or just to colistin, respectively.

Plasmid stability was determined incubating cells through successive growth cycles under non selective conditions. Every cycle was initiated by diluting (10^−2^) a stationary-phase liquid culture with fresh LB medium, and incubating at 37°C with gently shaking (200 rpm) during 16-20 h.

Plasmid curing was performed by using ethidium bromide in a gradient (g/L) of 0.05, 0.1 and 0.2, a 10^−3^ dilution of a stationary-phase liquid culture with fresh LB medium, incubating at 37°C with gently shaking (200 rpm) during 16-20 h and, after that, inoculating in parallel non-selective (LB) and colistin-selective (LB supplemented with 2 mg/L colistin) semi-solid media. Curing efficiency was determined by comparing the number of growing colonies and, as expected (Buckner et al., 2018), optimal efficiency (90%) was found by using 0.1 g/L of ethidium bromide, the maximal non-inhibitory concentration since no visible growth was observed with 0.2 g/L in liquid cultures.

### Quantitative PCR

Gene expression analyses were performed with cells growing in liquid cultures (Mueller-Hinton II). After overnight growth, cultures were renewed by diluting 1/10 with fresh media supplemented with 0.2% arabinose and incubated at 37°C with gently shaking (200 rpm). When cell cultures reached 0.3-0.5 OD_600nm_, they were quickly cooled on ice, centrifuged and processed for RNA extraction (Aurum Total RNA Minikit, Bio-Rad), reverse transcription (PrimeScript™ RT reagent Kit, Takara) and to remove genomic DNA (TURBO DNA-free kit, Ambion), according to the manufacturer’s protocols. SYBRgreen real-time quantitative assays were made by using the SYBR^®^ Premix Ex Taq™ II (Tli RNaseH Plus; Takara Bion Inc.) and an Applied Biosystems® Step One PCR System. Oligo Primer Analysis Software v. 7 was utilized to design primer sequences with optimal amplification efficiencies (supplementary Table II). The normalized relative quantities (NRQ) of transcripts were obtained by the 2^−ΔΔCt^ calculation method with the expression of *recA* gene used as calibration reference, and the *E. coli* strain ATCC25922 grown in the same conditions as reference for normalization (basal conditions), with every experimental condition including three technical replicates (triplicated reactions in the same qPCR) and two biological replicates from fully independent experiments. Ratios between the mean NRQ for every treatment and control conditions and the standard error of the ratios were calculated according to previously reported methods (Rieu et al., 2009).

### Pulsed Field Gel Electrophoresis

Determination of the genetic relationships among *E. coli* isolates was performed by macrorestriction with XbaI followed by PFGE (Chef DRII, Bio-Rad), according to the PulseNet protocol with pulses oscillating from 2.16 to 63.8 s for 21.5 hours (Ribot et al., 2006) with *S*. Braenderup used as molecular weight standard and the gel was stained with ethidium bromide. Plasmid composition analysis was performed by PFGE in the same conditions described above, after incubation of plugs with S1 nuclease (Thermo Fisher Scientific) according to the manufacturer recommendations. For plasmid hybridization, the PFGE-S1 was transferred to a nylon membrane and hybridized to a digoxigenin-labelled probe amplified from the *Kluyvera*-like *arnB* gene detected in ZTA15/00213-1EB1 strain, by using specific primers (supplementary Table II) according to the manufacturer instructions (Sigma-Aldrich).

### Genome analysis

DNA were extracted by QIAGEN DNeasy Blood and Tissue Kit and sequencing libraries were prepared using the Nextera XT kit and sequenced on a MiSeq (Illumina) using v3 reagents with 2 × 300 cycles. Isolates ZTA15/00213-1EB1 and ZTA15/00702EC produced 493612 and 473653 reads, respectively, which were assembled using SPAdes v 3.9.0. The draft genomes were 5076949 and 5092360 bp lengths, were composed by 452 and 449 contigs with N50 values of 28806 and 20026, respectively, corresponding to 20X coverage in both cases. WGS drafts of both genomes have been deposited in GenBank as a BioProject with ID PRJNA647041 (accessions JACEFD000000000 and JACEFE000000000, respectively). pArnT1 sequence (60178 bp) was deduced from JACEFD000000000 by assembling the contigs NODE_75 (24898) bp, NODE_304 (595 bp) and NODE_51 (34939 bp), in this order. The joining through NODE_304 was confirmed by PCR with primers pArnT1a (supplementary table II).

## Results

### Colistin resistance determinants of E. coli from farm animals

Colistin resistant isolates of *E. coli* were selected around Spain from feces of healthy animals of three major species targeted for food production; swine, poultry (turkey) and bovine (Table 1 and supplementary Table I). The *mcr-1* gene was found in 69 out of the 74 isolates, followed by *mcr-4* in two strains among which one also shared *mcr-1*. Besides *mcr-1* or *mcr-4* genes, we identified fifteen isolates with missense-polymorphisms in the *pmrAB* operon that were not found in *E. coli* K12 or ATCC 25922 nor in colistin susceptible strains recently isolated from farm animals (supplementary Table I; Quesada et al., 2015). The genotype of 4 isolates remained lacking any candidate for resistance determinant.

**Table 1.**
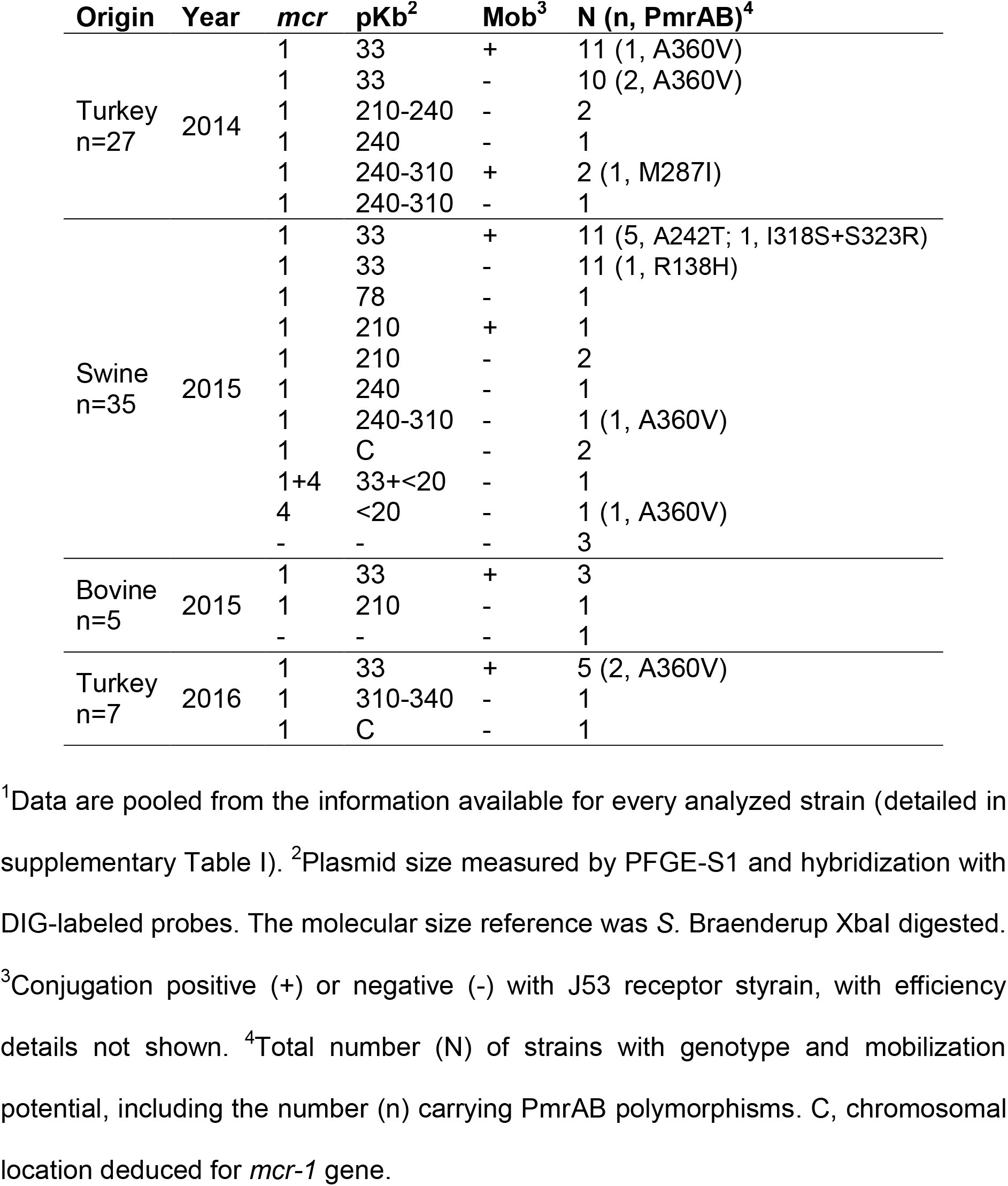
Colistin resistance determinants in *E. coli* from healthy animals^1^.

### Mobilization potential of colistin resistance determinants

PFGE-S1 and hybridization to a specific probe showed that 66 out of the 69 *mcr-1* genes detected in this work are located in plasmids which sizes are near 33Kb, 78Kb or higher than 200Kb (Table 1 and supplementary Table I), fitting almost unambiguously the ranges covered respectively by IncX4, IncI2 or IncHI2 replicons that carried more than 90% of *mcr-1* genes in previously analyzed enterobacteria (Matamoros et al., 2017). Besides, the two *mcr-4* genes hybridized to a plasmid lower than 20Kb that would correspond to the low molecular weight (8.7Kb) ColE1 replicon previously linked to this resistance determinant (Carattoli et al., 2017).

Conjugation assays showed that like the four strains lacking known colistin resistance determinants, none of the three strains that share chromosomally located *mcr-1* genes, the two ColE10-like plasmids with *mcr-4*, nor the unique plasmid carrying *mcr-1* in the size range o+f an IncI2 replicon, mobilized their phenotype to the recipient strain (Table 1 and supplementary Table I). In contrast, the efficiency of plasmids carrying *mcr-1* determinants to transfer colistin resistance by conjugation (47.8%) was much higher for IncX4-like (57.7%) than IncHI2-like (7.2%), the two major plasmids detected, in 52 and 13 isolates, respectively.

### Signaling of colistin resistance on the expression of arnBCADTEF and eptApmrAB operons

We measured the mRNA expression of *arnBCADTEF* and *eptApmrAB* operons from isolates representing the different determinants identified, *mcr-1* and/or *mcr-4* and the five polymorphisms found in PmrAB; R138H in PmrA and A360V, A242T, I318S+S323R and M287I in PmrB, plus all those which resistance determinant(s) remain(s) unknown (Figure 2). Accumulations of mRNA from both operons, that were detected by probing *arnB* and *eptA*, the first coding sequence of every one, was strongly down-regulated in strains presenting chromosomal *mcr-1*, IncX4-like carrying *mcr-1* and/or ColE10-like *mcr-4*, independently of polymorphism detected in PmrAB. In contrast, the two previously characterized swine isolates ZTA11/01748EC and ZTA13/02182EC that present *mcr-1* besides polymorphisms PmrB-V161G or PmrA-S39I+R81S, respectively (Quesada et al., 2015; 2016), mutations that had been shown to be involved in colistin resistance (Sun et al., 2009), shared *arnB* expression up-regulated (Figure 2). Thus: i) *mcr* expression negatively signals mRNA accumulation from both operons; ii) as previously shown, PmrAB mutations that self-activate it constitutively and confer colistin resistance up regulate *arnB* and, to a lower extent, *eptA* expression (Sun et al., 2009); and iii) up regulation by PmrAB mutations that confer colistin resistance dominates over down regulation by *mcr* genes in strains carrying both resistance determinants. Therefore, since arnB/eptA remained repress in isolates carrying *mcr* genes and PmrAB polymorphisms detected in this work, these might not be functionally relevant for colistin resistance. In addition, the four isolates lacking known resistance determinants share both operons repressed, like the strains expressing *mcr* genes (Figure 2).

**Figure 2.**
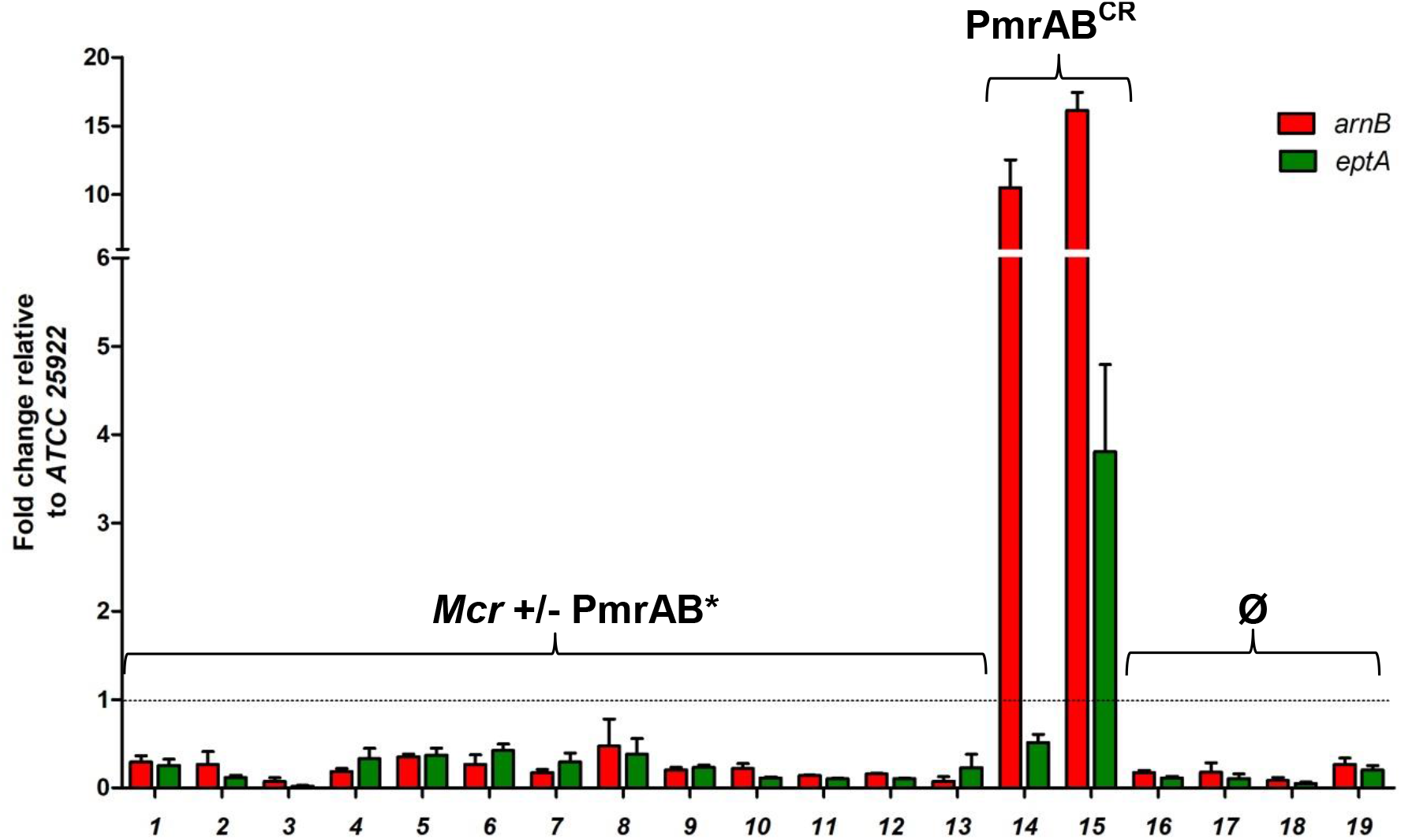
Signaling of colistin resistance determinants on *arnBCADTEF* and *eptApmrAB* operons expression. Liquid cultures of indicated strains were incubated until mid-exponential phase of growth, and processed for RNA extraction, cDNA synthesis and qPCR to quantify *arnB* and *eptA* transcript accumulation, by using *recA* as an endogenous calibrator and the *E. coli* ATCC25922 strain for normalization of gene expression, which level is indicated by a discontinuous line. Strains and genotypes (*mcr* genes and PmrAB polymorphisms, supplementary Table I) are: 1, ZTA15/02267EB1 (chromosomal *mcr-1*); 2, ZTA15/00389EB1 (*mcr-1*); 3, ZTA15/01072COL1 (*mcr-1* plus A242T-PmrB); 4, ZTA15/01844EB1 (*mcr-1* plus A242T-PmrB); 5, ZTA15/02333EB2 (*mcr-1* plus A242T-PmrB); 6, ZTA15/01825EB1 (*mcr-1* plus A242T-PmrB); 7, ZTA15/00233EB1 (*mcr-1* plus A242T-PmrB); 8, ZTA15/00750EB1 (*mcr-4* + A360V-PmrB); 9, ZTA15/00630EB1 (*mcr-1* + A360V-PmrB); 10, ZTA15/01313EB1 (*mcr-1* + I318S,S323R-PmrB); 11, ZTA14/00808EB (*mcr-1* + M287I-PmrB); 12, ZTA15/01196EB1 (*mcr-1* + R138H-PmrA); 13, ZTA15/00420EB1 (*mcr-1* + *mcr-4*); 14, ZTA11/01748EC (*mcr-1* + V161G-PmrB, Quesada et al., 2016); 15, ZTA13/02182EC (*mcr-1* + S39I,R81S-PmrA, Quesada et al., 2016); 16, ZTA15/00213-1EB1 (unknown); 17, ZTA15/01360EB1 (unknown); 18, ZTA15/00235EB1 (unknown); 19, ZTA15/00702EC (unknown). PmrAB* indicates strains with PmrAB polymorphisms detected in this work, whereas PmrAB^CR^ are already known to confer colistin resistance (Sun et al 2009).

### Finding of a Kluyvera-like arnBCADTEFT operon in colistin resistant E. coli

PFGE analysis revealed that ZTA15/00213-1EB1, ZTA15/01360EB1 and ZTA15/00235EB1, three out of the four isolates remaining without known determinant for colistin resistance, are closely related (Figure 3A) and share two plasmids, roughly co-migrating with the 55 Kb and 105 Kb bands of the size standard, whereas the remaining isolate ZTA15/00702EC presented a different profile (Figure 2B).

**Figure 3.**
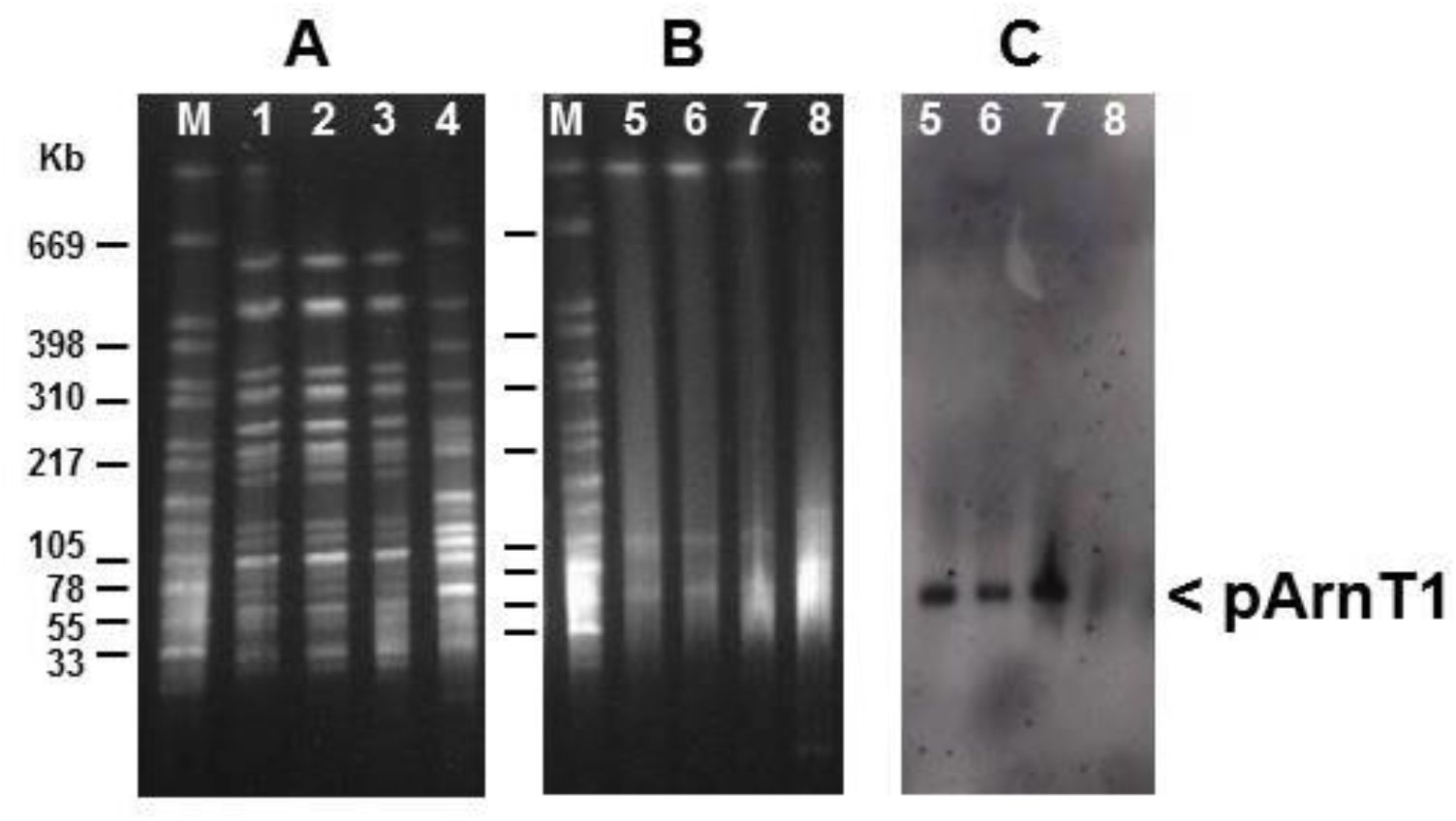
PFGE analysis of strains lacking previously known colistin resistance determinants. A, XbaI macrorestriction and genomic profiling; B, Nuclease S1 and plasmid profiling; C, Southern blot of PFGE-S1 hybridized to a DIG-labelled probe of the *arnB* sequence, *Kluyvera*-like, PCR amplified from strain ZTA15/00213-1EB1. 1, ZTA15/00213-1EB1; 2, ZTA15/01360EB1; 3, ZTA15/00235EB1; 4, ZTA15/00702EC.

Genome sequencing was performed for isolates ZTA15/00213-1EB1 and ZTA15/00702EC representing the two strains with different PFGE types identified, which were isolated from bovine and porcine, respectively (supplementary Table I). In addition to the search already performed by PCR of *mcr* genes and sequencing PmrAB coding sequences, both genomes were screened for *mcr-1* to *mcr-10* genes (Wang et al., 2010) and polymorphisms in *phoPQ*, *mgrB* or *pmrD* genes (Olaitan et al., 2014), without finding any candidate that might confer colistin resistance, although other antibiotic resistance determinants were detected (supplementary Table III). However, searching the genes involved in LPS modification and looking for their possible structural alterations, we found a non-native *arnBCADTEF* operon of 7382 bp length in isolate ZTA15/00213-1EB1, in addition to its *E. coli*-like counterpart of 7351 bp (supplementary Table IV). The foreign *arnBCADTEF* operon gives the best match (97.7%) by BLASTN with *Kluyvera ascorbata* OT2 genome found in the RefSeq Genome Database (refseq_genomes) available at NCBI. The identity between *arnBCADTEF* DNA sequences from *K. ascorbata* and the *Kluyvera*-like operon found in ZTA15/00213-1EB1 is above 96% along their seven coding sequences, although it is even higher (above 99%) among *E. coli*-like sequences from its own operon and that of *E. coli* K12 or from ZTA15/00702EC, the other isolate lacking previously known colistin resistance determinants (supplementary Table IV). The identities between *Kluyvera*-like and *E. coli*-like *arnBCADTEF* sequences decreased to 53-74%, still closely related but far-below the hybridization threshold that determines specificity of primers (see above) and DNA probe (next section) used in this work. Besides that, the seven coding sequence of *arnBCADTEF* operons found in the genomes of the two *E. coli* isolates are complete and (apparently) fully functional (supplementary Table IV).

### pArnT1, an IncFII plasmid carrying a Kluyvera-like arnBCADTEF operon

The *Kuyvera*-like *arnBCADTEF* operon from ZTA15/00213-1EB1 genome is located in a 60.1 Kb contig spanning 79 putative coding sequences, many of which present plasmidic features like *tra* and *trb* genes for mobilization functions and *repA1* for replication initiation belonging to the IncFII replicon-type (Carattoli et al., 2014). In addition, it presents three antimicrobial resistance genes; *bla*-TEM, *sul1* and *emrE*, for a class-A β-lactamase, a dihydropteroate synthase and a multidrug efflux protein, respectively. Most of the remaining 53 Kb DNA surrounding the *Kluyvera*-like operon is closely related with fragments from two previously described replicons (Figure 4). This plasmid will be named hereafter pArnT1.

**Figure 4.**
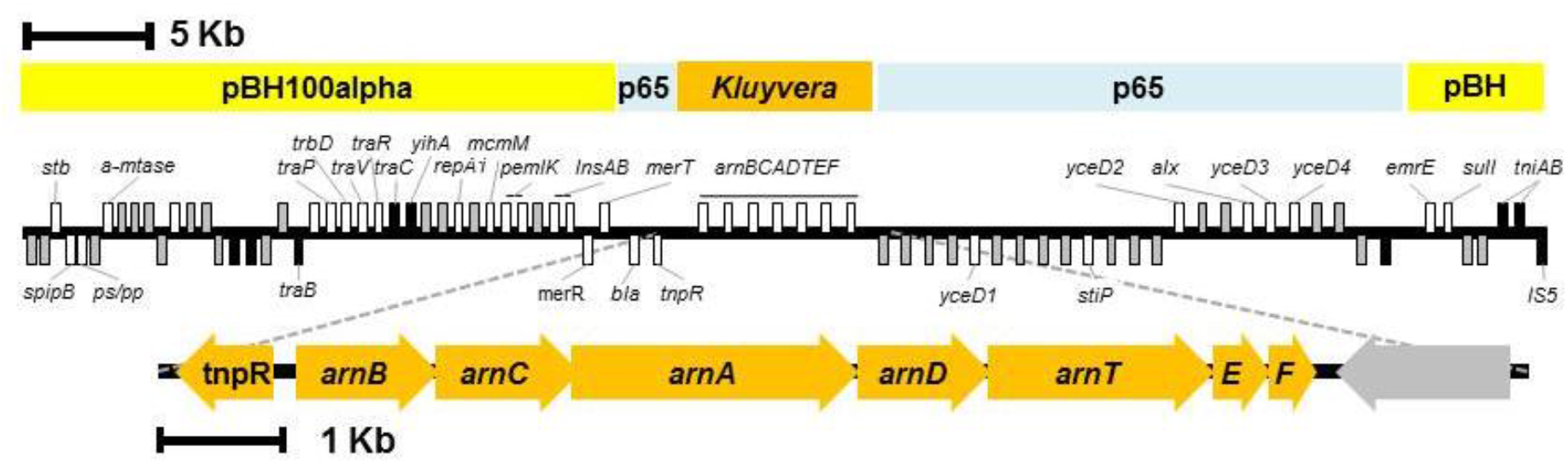
Structure of pArnT1. Plasmid sequence is covered over almost its entire length with discontinuous regions sharing high identity with *E. coli* plasmids pBH100alpha (CP025254.1) and p65 (MT077888.1), in addition to the *arnBCADTEF* operon sequence from *K. ascorbata* OT2 (NZ_RHFN01000006.1). TBLASTN identified 78 coding sequences, which are represented by boxes above or below for sense or antisense strands, respectively. Black boxes correspond to truncated sequences and grey boxes to hypothetical proteins lacking known function.

The presence of pArnT1 in colistin resistant strains of *E. coli* was evidenced by PFGE-S1 and Southern hybridization to a *Kluyvera*-like *arnB* probe (Figure 2C). Isolates ZTA15/00213-1EB1, ZTA15/00235EB1 and ZTA15/01360EB1, which share PFGE profiles (Figure 2A and B), presented a plasmid migrating just above 55 Kb that fits the 60.1 Kb of pArnT1 sequence and that specifically hybridized to the *Kluyvera*-like *arnB* probe.

### pArnT1 stability and Kluyvera-like arnBCADTEF expression are associated to colistin resistance

The functionality of pArnT1 could not be addressed from its horizontal mobilization since neither the conjugation (see above) nor the transformation (not shown) of the colistin resistance phenotype from isolates lacking previously known determinants were successful. The opposite evidence, a loss of pArnT1 that correlates with reduction of colistin resistance, was approached. Stability of the phenotype was different among the three strains sharing pArnT1 (Figure 5). Whereas isolates ZTA15/00213-1EB1 and ZTA15/00235EB1 steadily preserved colistin resistance and pArnT1 after (up to four complete cycles of) growth in media lacking colistin, ZTA15/01360EB1 lost both traits without selection and only one clone out of 5 grown on colistin media retained the *arnB-Kluyvera* like sequence (Figure 5A). Accordingly, mRNA expression from the plasmid carried *arnBCADTEF* operon was detected in isolates ZTA15/00213-1EB1 and ZTA15/00235EB1 but not in ZTA15/01360EB1 (Figure 5B), which *pmrAB* coding sequences were amplified from the clone growing on colistin (Figure 5A) finding out the V161G polymorphism of PmrB (not shown), a mutation that constitutively activates it and confers colistin resistance by itself (Olaitan et al., 2014) and that must have occurred during in vitro growth because originally was not present (supplementary Table I). Thus, mRNA expression from the *Kluyvera*-like *arnBCADTEF* operon is linked to colistin resistance, although its DNA sequence might become inactivated like that of ZTA15/01360EB1 isolate.

**Figure 5.**
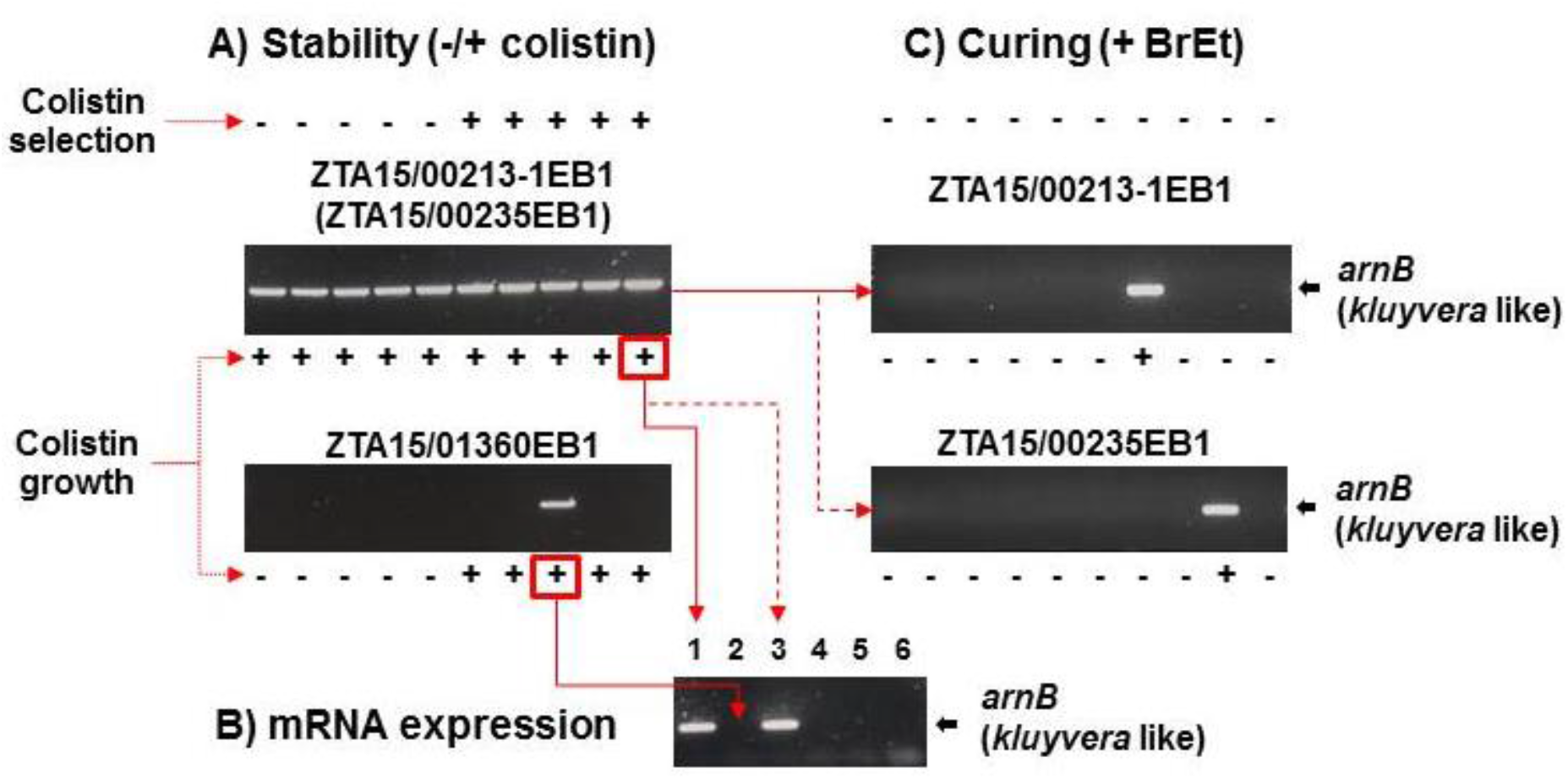
Association of the *Kluyvera*-like *arnBCADTEF* operon to colistin resistance. **A.** The three *E. coli* isolates sharing the pArnT1 plasmid; ZTA15/00213-1EB1, ZTA15/00235EB1 and ZTA15/01360EB1, were grown overnight with or without antibiotic selection (colistin, 2 mg/L). Five clones were selected from every strain and condition, their growth in colistin was tested and, after DNA purification, the *Kluyvera*-like *arnB* sequence was detected by PCR. Identical results were obtained for isolates ZTA15/00213-1EB1 and ZTA15/00235EB1 (not shown). **B.** RNA was isolated from one clone selected on colistin selective media from every strain and, after cDNA synthesis, the DNA fragment spanning *arnB* gene was amplified by RT-PCR with primers indicated in supplementary Table II. 1, ZTA15/00213-1EB1; 2, ZTA15/01360EB1; 3, ZTA15/00235EB1; 4, ZTA15/00702EC; 5, *E. coli* ATCC25922; 6, negative control with water instead cDNA. **C.** The plasmid was cured by growing cells overnight with ethidium bromide (100 mg/L). Ten colonies were selected on non-selective media from every strain, their growth on colistin was tested and, after DNA purification, the *Kluyvera*-like *arnB* sequence was detected by PCR.

A functional proof for the involvement of pArnT1 in colistin resistance was reached by plasmid curing. After one growth cycle in media containing a sub-inhibitory concentration of ethidium bromide, the two independent isolates that stably preserve and express the *Kluyvera*-like *arnBCADTEF* operon without colistin selection, ZTA15/00213-1EB1 and ZTA15/00235EB1, loss in parallel the ability to grow on colistin selective media and pArnT1, tagged by PCR amplification of the *Kluyvera*-like *arnB* coding sequence (Figure 5C). In contrast, the colistin resistance trait that remained in 10% of cells after plasmid curing was closely linked to pArnT1.

## Discussion

The two acquired mechanisms conferring colistin resistance to enterobacteria, chromosomal mutations that constitutively activate PmrAB and plasmid-mobilized *mcr* genes, promote opposite signaling on mRNA expression from *arnBCADTEF* and *eptApmrAB* operons, encoding key enzymes for LPS modification. This work evidences that polymyxin-resistant enterobacteria lacking previously known determinants show *mcr*-like down-regulation of target genes, evoking structural modification of their lipid A. We identified pArnT1, a new plasmid carrying a *Kluyvera*-like *arnBCADTEF* operon, in this category of colistin-resistant *E. coli* and found associations between element functioning and antibiotic susceptibility.

The analysis of *E. coli* from feces of healthy animals performed in this work (Table 1 and supplementary Table I) showed that *mcr-1* was the main determinant for colistin resistance in this environment at the time of screening (2014-2016). *mcr-1* was carried by different plasmids among which predominated IncX4 (70%), accurately identified like a 33 Kb element by PFGE-S1 plus specific hybridization. These IncX4-like elements were highly mobilizable (44%), in contrast to the other *mcr* gene identified in this work, *mcr-4*, found linked to a low molecular weight ColE1-like plasmid and that was not transferable in conjugation essays (Table 1 and supplementary Table I), like the mega-plasmid carrying *mcr-3* gene previously described (Hernandez et al., 2017).

Among the 10 *mcr* genes described so far in Gram-negative bacteria, *mcr-1* was the first plasmidic colistin-resistant determinant identified (Liu et al., 2016) and, not casually, it is the commonest and worldwide spread and largely predominates in *E. coli* (Kheder et al., 2020). The health status of animals from where *E. coli* is isolated seems not conditioning the prevalence of *mcr-1*, equally high (>90%) in China than in Spain from healthy poultry and swine (Huang et al., 2017, Table 1) or in France from diseased pigs (Delanoy et al., 2017), among which roughly 50% were mobilized by conjugation, similarly to this work (Table 1). However, local outbreaks may explain that *E. coli* from post-weaning diarrhea in Spain presented a higher prevalence of *mcr-4* over *mcr-1* or *mcr-5*, the other two determinants frequently found (Garcia et al., 2018).

Besides *mcr* genes, chromosomal mutations and other(s) yet unknown acquired determinant(s) are systematically detected among colistin resistance strains, although their prevalence is variable depending on the environment from which *E. coli* is isolated. Thus, colistin resistant *E. coli* from human inpatients in Paris region presented low carriage of *mcr* genes (5%, *mcr*-1 uniquely found), a high rate of missense polymorphisms in *pmrA* or *pmrB* coding sequences (69%) and a significant fraction (26%) lacking known resistance determinant (Bourrel et al., 2019). In contrast, a low fraction (4%) of isolates remained without known determinants from diseased animals analyzed in Spain (Garcia et al., 2018; Garcia et al., 2019), fitting the prevalence found in this work (Table 1). In our screening we sampled bacteria from healthy animals and we found 15 isolates presenting chromosomal mutations in PmrAB that cannot be detected in colistin susceptible strains, a first clue of their possible involvement in colistin resistance (Quesada et al., 2015). However, neither of these resulted in the up regulation of *eptA* and/or *arnB* expression, in contrast to previously known mutations that activate PmrAB constitutively (Sun et al., 2009; Figure 2), and might not be considered true determinants for colistin resistance.

A significant fraction of isolates analyzed in this work (5.4%) lacked previously known acquired determinants for colistin resistance and most of them carry pArnT1, a new 60.1 Kb plasmid belonging to the IncFII replicon type. This element is a chimera among fragments from two replicons already described, pBH11alpha (CP025254.1) and p65 (MT077888.1), and a genomic fragment closely related (97.7% identity) to the *arnBCADTEF* operon from the opportunistic human pathogen *K. ascorbata*. Since, to our knowledge, plasmid mobilization of *arnBCADTEF* operon has been never found before, its occurrence in *E. coli* isolates sharing low susceptibility to colistin, lacking previously known resistance determinants and presenting down-regulated the expression from their own native *arnBCADTEF* operon, in addition to the linkage found by plasmid curing between phenotype and genotype, strongly suggests a role for pArnT1 in colistin resistance. However, although repeatedly assayed, conjugation of this element by using the same essay that evidenced the efficient transfer of roughly 50% of plasmids carrying *mcr-1* genes was unsuccessful. Indeed, pArnT1 presents truncated coding sequences for two transfer functions, TraB and TraC (Figure 4), which probably explains that element spread in the analyzed environment is limited to vertical transmission of clonally related isolates (Figure 3A).

The *Kluyvera*-like *arnBCADTEF* operon found in pArnT1 is apparently functional and, like its *E. coli* orthologs (Table 2), might be related with LPS modification of enterobacteria. Although little is known about colistin resistance mechanisms of *K. ascorbata*, a low susceptible strain presented the *mcr-1* determinant mobilized by an IncI2 replicon (Zhao et al., 2016), which sequence is not related to pArnT1 (not shown).

In contrast to *mcr* genes, several of which (*mcr-1, −2, −3, −4* and −*5*) have been proved to confer colistin resistance upon cloned in *E. coli* and transcribed from expression vectors, little information is available about any similar approach focused on the *arnBCADTEF* operon, despite parallel roles of their encoded functions on LPS modification (Figure 1). Expression in *E. coli* of its own *arnT* coding sequence from the same pBAD vector used to express *mcr* genes only increased marginally colistin resistance (Zhang et al., 2019). However, *arnT* coding sequence was expressed lacking the other six gene functions carried by the *arnBCADTEF* operon plus the unlinked *ugd* gene, all together providing L-Ara4N-undecaprenyl phosphate, the substrate for ArnT activity (Figure 1). Future works are required to address the heterologous and controlled expression of *Kluyvera*-like or *E. coli*-like *arnBCADTEF* operons, an approach that would be required to decipher the biochemical and physiological consequences derived from the functioning of these elements.

Presented evidences suggest that enterobacteria have gained a new colistin resistance determinant by plasmid mobilization of genes for modification of lipid A with L-Ara4N, in addition to *mcr* genes that perform it by PEtN addition. pArnT1 or similar elements should be screened in enterobacteria, mainly in those lacking previously known colistin resistance determinants, to know their prevalence in animal and human environments, evaluating the risk of this new challenge from a One Health perspective.

## Acknowledgements

M.R.I. and P.M.V. received PhD fellowships, respectively, from the “Fundación Tatiana de Guzman El Bueno” (Spain), and the FPI Program (BES-2017-080264) from the Spanish Ministry of Science, Innovation and Universities. This work has been funded by the Spanish Ministry of Economy, Industry and Competitiveness (MINECO, actually MICINN, Grant AGL2016-74882-C3), and the Junta de Extremadura and FEDER (IB16073 and GR15075) of Spain.

---

**Supplementary Table I.**
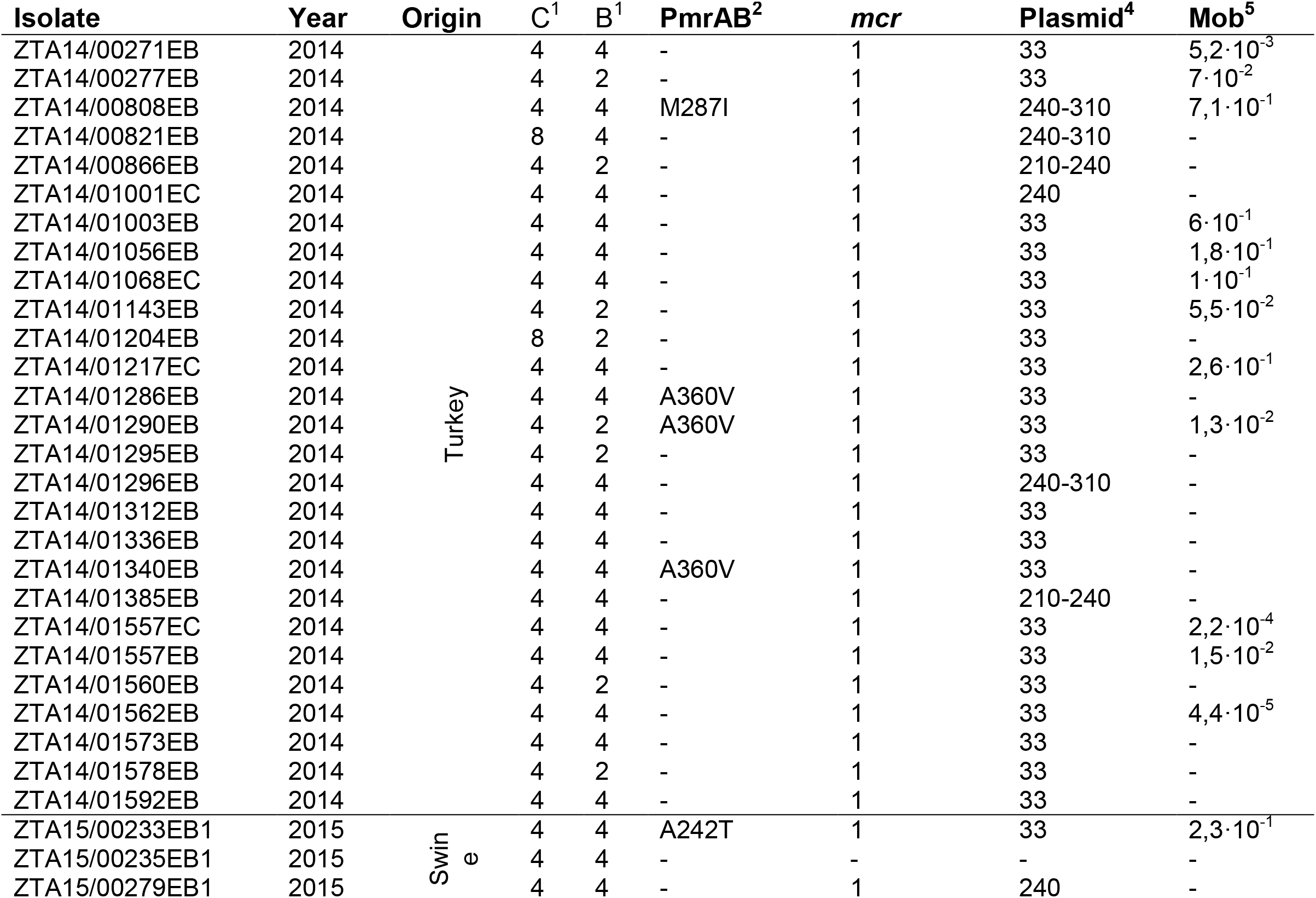
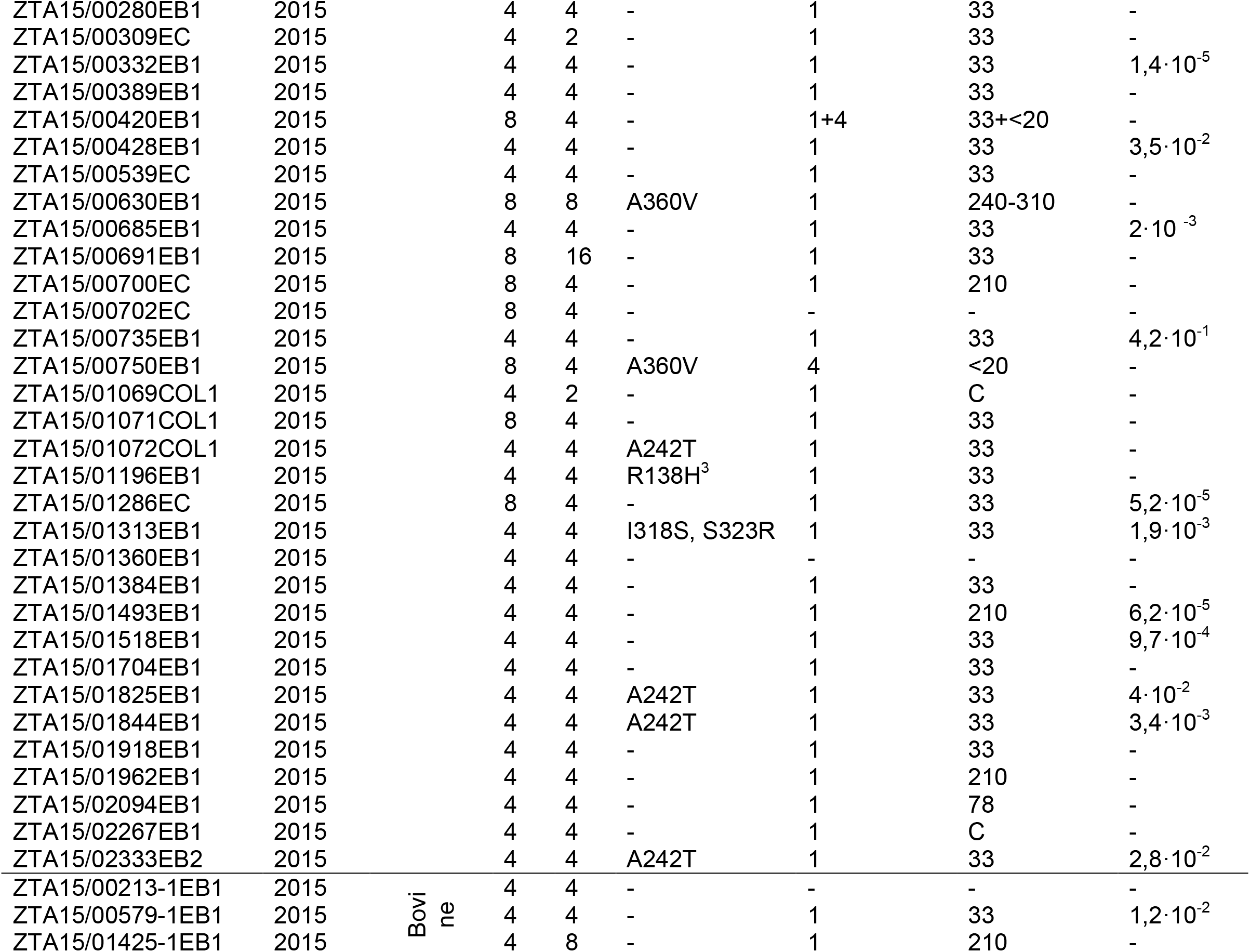
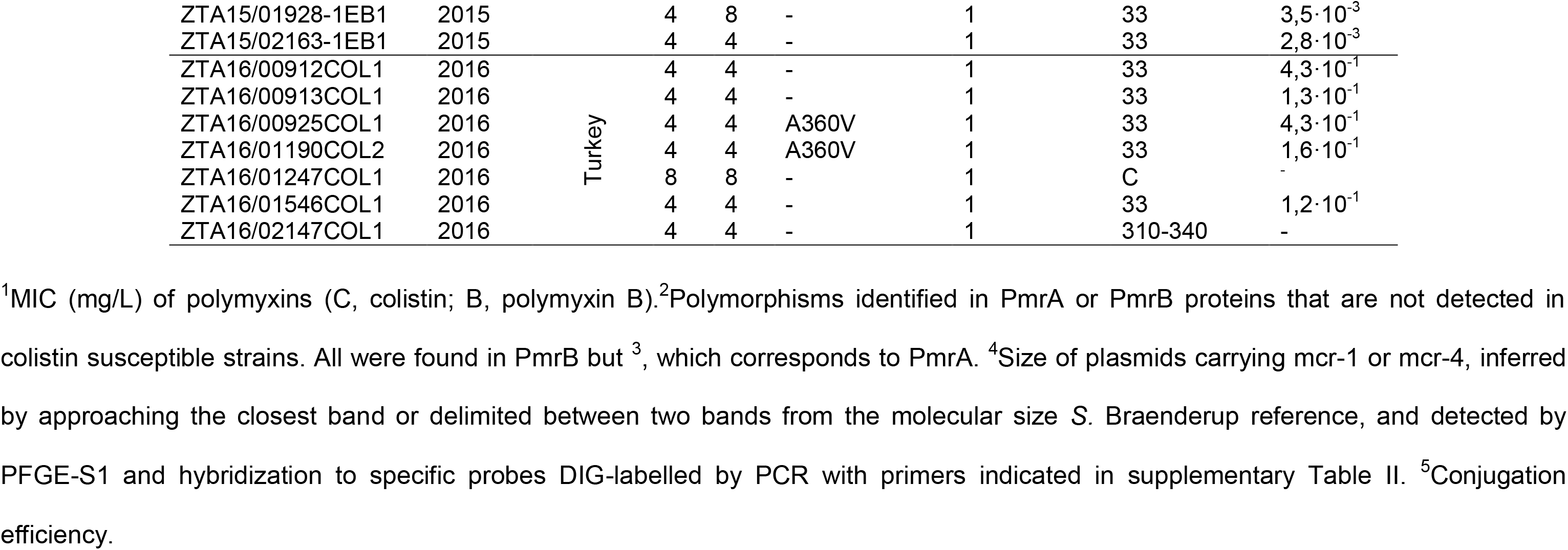
Phenotype and genotype analysis of colistin resistant *E. coli* strains from healthy animals.

**Supplementary Table II.**
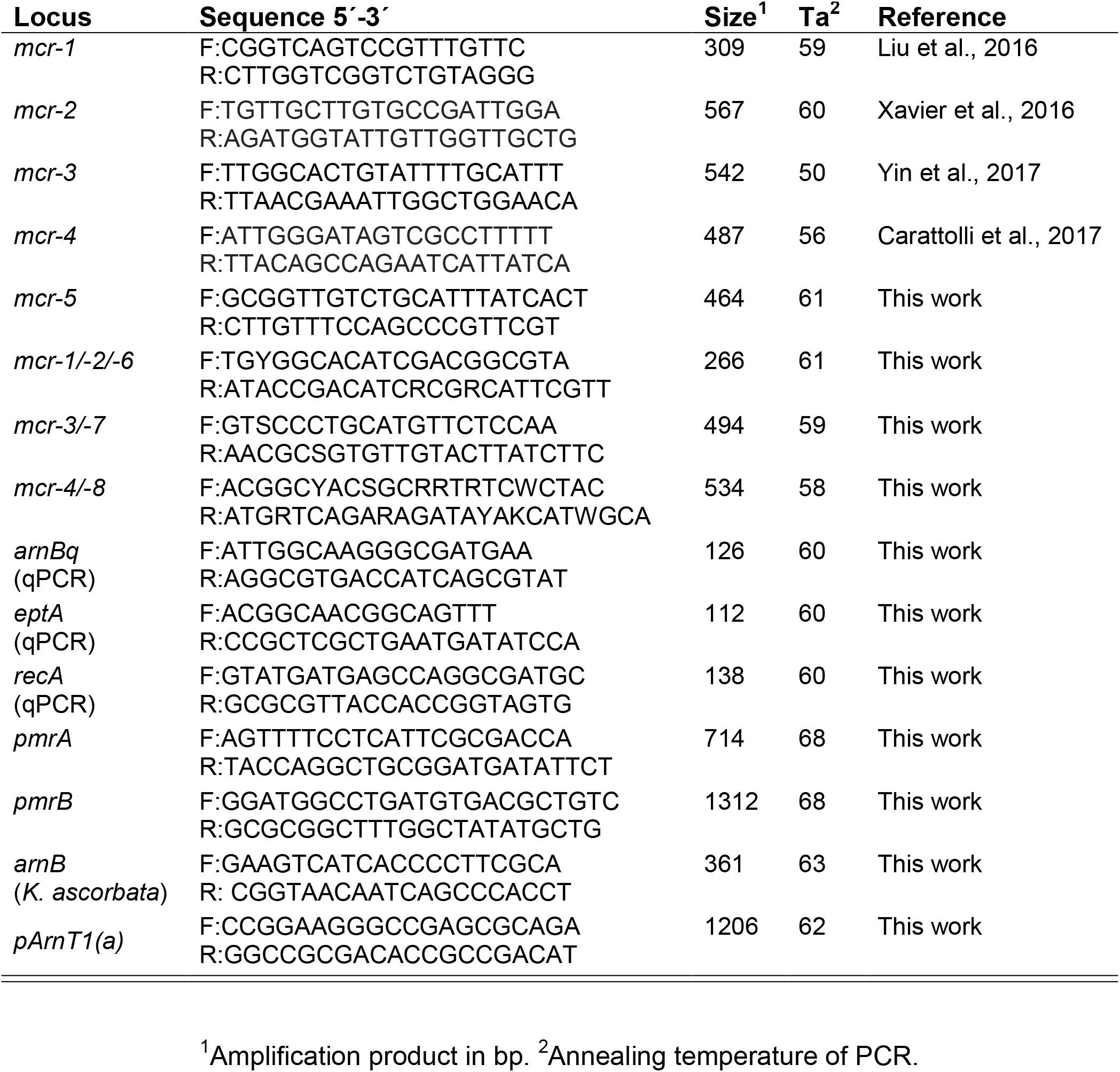
Primers used in this work.

**Supplementary Table III.** Genome analysis of colistin resistant isolates^1^.

|  | **loci** | **ZTA15/00213-1EB1** | **ZTA15/00702EC** |
| --- | --- | --- | --- |
| **Antimicrobial resistance** | *aad1* | 100 | 100 |
|  | *aph(3')-1a* | 99.9 | - |
|  | *bla-SHV-12* | 100 | - |
|  | *bla-TEM1b* | - | 100 |
|  | *bla-TEM-150* | 100 | - |
|  | *cmlA1* | 99.9 | - |
|  | *dfrA1* | - | 100 |
|  | *ermB* | 99.9 | - |
|  | *mdf(A)* | 98.5 | 99.8 |
|  | *sul1* | 100 | 100 |
|  | *sul3* | 100 | - |
|  | *tetA* | - | 100 |
|  | *tetM* | 96.2 | - |
| **Replicons** | ColpVC | - | 99.0 |
|  | IncFIB<br>(AP001918) | - | 99.0 |
|  | IncFIB (H89-PP) | - | 99.0 |
|  | IncFII | 100 | 100 |
| | IncI1-($\gamma$) | 99.3 | - |
|  | IncX1 | 98.7 | 98.7 |
|  | IncY | - | 99.1 |
|  | p0111 | 99.2 | - |

<sup>1</sup>Identities (%) between loci detected by WGS analysis with ResFinder 3.2 (Zankari et al., 2013) and PlasmidFinder 2.0 (Caratolli et al., 2014).

**Supplementary Table IV ^1^.** Conservation between *arnBCADTEF* operons.

| Loci |  | size | K213 | Kas | E213 | E702 |
| --- | --- | --- | --- | --- | --- | --- |
| *arnBCADTEF* | Kas | 7382 | 97.70 |  |  |  |
|  | E213 | 7351 | 68.80 | 68.74 |  |  |
|  | E702 | = | 68.78 | 68.80 | 99.31 |  |
|  | K12 | = | 68.83 | 68.81 | 99.27 | 99.93 |
| *arnB* | Kas | 1137 | 98.95 |  |  |  |
|  | E213 | = | 70.96 | 71.05 |  |  |
|  | E702 | = | 70.88 | 70.96 | 99.74 |  |
|  | K12 | = | = | = | = | 100 |
| *arnC* | Kas | 978 | 97.15 |  |  |  |
|  | E213 | 966 | 74.41 | 74.61 |  |  |
|  | E702 | = | = | 74.82 | 99.69 |  |
|  | K12 | = | = | = | = | 100 |
| *arnA* | Kas | 1983 | 97.33 |  |  |  |
|  | E213 | 1980 | 71.66 | 71.71 |  |  |
|  | E702 | = | = | 71.86 | 97.88 |  |
|  | K12 | = | = | = | = | 100 |
| *arnD* | Kas | 900 | 98.45 |  |  |  |
|  | E213 | 888 | 66.89 | 67.34 |  |  |
|  | E702 | = | = | = | 100 |  |
|  | K12 | = | = | = | = | 100 |
| *arnT* | Kas | 1653 | 97.28 |  |  |  |
|  | E213 | 1650 | 65.46 | 64.91 |  |  |
|  | E702 | = | = | = | 99.94 |  |
|  | K12 | = | = | = | = | 100 |
| *arnE* | Kas | 336 | 96.17 |  |  |  |
|  | E213 | 333 | 65.18 | 65.48 |  |  |
|  | E702 | = | = | = | 100 |  |
|  | K12 | = | 65.77 | = | 99.4 | 100 |
| *arnF* | Kas | 387 | 98.72 |  |  |  |
|  | E213 | 384 | 54.26 | 53.75 |  |  |
|  | E702 | = | = | = | 100 |  |
|  | K12 | = | 54.52 | 54.01 | 99.22 | 100 |

<sup>1</sup>*arnBCADTEF* sequences: K213, *E. coli* ZTA15/00213-1EB1(*Kluyvera*-like); Kas, *K. ascorbata* OT2 KA\_6; E213, *E. coli* ZTA15/00213-1EB1(*E. coli*-like); E702, *E. coli* ZTA15/00702EC (*E. coli*-like); K12, *E. coli* K-12 MG1655. Comparisons were performed by Clustal Omega (https://www.ebi.ac.uk/), from which identities (%) between sequence pairs are indicated. The equal symbol (=) denotes identical similarity than the above value.

## References

Buckner MMC, Ciusa ML, Piddock LJV. Strategies to combat antimicrobial resistance: anti-plasmid and plasmid curing. FEMS Microbiol Rev 2018; 42: 781–804.

Bourrel AS, Poirel L, Royer G, et al. Colistin resistance in Parisian inpatient faecal Escherichia coli as the result of two distinct evolutionary pathways. J Antimicrob Chemother 2019; 74: 1521–30.

Carattoli A, Villa L, Feudi C, Curcio L, Orsini S, Luppi A, Pezzotti G, Magistrali CF. Novel plasmid-mediated colistin resistance *mcr-4* gene in *Salmonella* and *Escherichia coli*, Italy 2013, Spain and Belgium, 2015 to 2016. Eurosurveill 2017; 22: 30589.

Carattoli A, Zankari E, García-Fernández A, Voldby Larsen M, Lund O, Villa L, Møller Aarestrup F, Hasman H. In silico detection and typing of plasmids using PlasmidFinder and plasmid multilocus sequence typing. Antimicrob Agents Chemother 2014; 58: 3895–903.

Delannoy S, Le Devendec L, Jouy E, Fach P, Drider D, Kempf I. Characterization of Colistin-Resistant *Escherichia coli* Isolated from Diseased Pigs in France. Front Microbiol 2017; 8: 2278.

García V, García-Meniño I, Mora A, Flament-Simon SC, Diaz-Jimenez -D, Blanco JE, Alonso MP, Blanco J. Co-occurrence of *mcr-1*, *mcr-4* and *mcr-5* genes in multidrug-resistant ST10 Enterotoxigenic and Shiga toxin-producing *Escherichia coli* in Spain (2006-2017). Int J Antimicrob Agents 2018; 52: 104–8.

García-Meniño I, Díaz-Jiménez D, García V, Toro M, Flarment-Simon SC, Blanco J, Mora A. Genomic Characterization of Prevalent *mcr-1, mcr-4*, and *mcr-5 Escherichia coli* Within Swine Enteric Colibacillosis in Spain. Front Microbiol. 2019; 10: 2469.

Green MR Sambrook J. Molecular cloning: a laboratory manual, 4^th^ edition. Cold Spring Harbor Laboratory Press, Cold Spring Harbor, New York, 2012.

Hernández M, Iglesias MR, Rodríguez-Lázaro D, Gallardo A, Quijada N, Miguela-Villoldo P, Campos MJ, Píriz S, López-Orozco G, de Frutos C, Sáez JL, Ugarte-Ruiz M, Domínguez L, Quesada A. Co-occurrence of colistin-resistance genes *mcr-1* and *mcr-3* among multidrug-resistant *Escherichia coli* isolated from cattle, Spain, September 2015. EuroSurveill 2017; 22: 30586.

Huang X, Yu L, Chen X, Zhi X, Yao X, Liu Y, Wu S, Guo Z, Yi L, Zeng Z, Liu JH. High Prevalence of Colistin Resistance and *mcr-1* Gene in *Escherichia coli* Isolated from Food Animals in China. Front Microbiol. 2017; 8: 562.

Khedher MB, Baron SA, Riziki T, Ruimy R, Raoult D, Diene SM, Rolain JM. Massive analysis of 64,628 bacterial genomes to decipher water reservoir and origin of mobile colistin resistance genes: is there another role for these enzymes? Sci Rep 2020; 10: 5970.

Liu YY, Wang Y, Walsh TR, Yi LX, Zhang R, Spencer J, Doi Y, Tian G, Dong B, Huang X, Yu LF, Gu D, Ren H, Chen X, Lv L, He D, Zhou H, Liang Z, Liu JH, Shen J. Emergence of plasmid-mediated colistin resistance mechanism MCR-1 in animals and human beings in China: a microbiological and molecular biological study. Lancet Infect Dis 2016; 16: 161–8.

Matamoros S, van Hattem JM, Arcilla MS, Willemse N, Melles DC, Penders J, Vinh TN, Hoa NT, Bootsma MCJ, van Genderen PJ, Goorhuis A, Grobush M, Molhoek N, Lashof AMLO, Stobbeingh EE, Verbrugh HA, de Jong MD, Schulsz C. Global phylogenetic analysis of *Escherichia coli* and plasmids carrying the *mcr-1* gene indicates bacterial diversity but plasmid restriction. Sci Rep 2017; 7: 15364.

Olaitan AO, Morand S, Rolain JM. Mechanisms of polymyxin resistance: acquired and intrinsic resistance in bacteria. Front Microbiol 2014; 5: 643.

Quesada A, Ugarte-Ruiz M, Iglesias MR, Porreri MC, Martinez R, Florez-Cuadrado D, Campos, MJ, Garcia M, Piriz S, Dominguez L. Detection of plasmid mediated colistin resistance (MCR-1) in *Escherichia coli* and *Salmonella enterica* isolated from poultry and swine in Spain. Res Vet Sci 2016; 105: 134–5.

Quesada A, Porrero MC, Téllez S, Palomo G, García M, Domínguez L. Polymorphism of genes encoding PmrAB in colistin-resistant strains of *Escherichia coli* and *Salmonella enterica* isolated from poultry and swine. J Antimicrob Chemother 2015; 70: 71–4.

Raetz CR, Reynolds CM, Trent MS, Bishop RE. Lipid A modification systems in gram-negative bacteria. Annu Rev Biochem 2007; 76: 295–329.

Ribot EM, Fair M, Gautom R, Cameron D, Hunter S, Swaminathan B, Barrett TJ. Standardization of pulsed-field gel electrophoresis protocols for the subtyping of *Escherichia coli* O157: H7, *Salmonella*, and *Shigella* for PulseNet. Foodborne Path Dis 2006; 3: 59–67.

Rieu I, Powers SJ. Real-time quantitative RT-PCR: design, calculations, and statistics. Plant Cell 2009; 21: 1031–3.

Sun S, Negrea A, Rhen M, Andersson DI. Genetic analysis of colistin resistance in *Salmonella enterica* serovar Typhimurium. Antimicrob Agents Chemother 2009; 53: 2298–305.

Wang C, Feng Y, Liu L, Wei L, Kang M, Zong Z. Identification of novel mobile colistin resistance gene *mcr-10*. Emerg Microbes Infect 2020; 9: 508–16.

Xavier BB, Lammens C, Ruhal R, Kumar-Singh S, Butaye P, Goossens H, Malhotra-Kumar S. Identification of a novel plasmid-mediated colistin-resistance gene, *mcr-2*, in *Escherichia coli*, Belgium, June 2016. Euro Surveill. 2016; 21: 30280.

Yin W, Li Hui, Shen Y, Liu Z, Wang S, Shen Z, Zhang R, Walsh TR, Shen J, Wang Y, Bush K. Novel Plasmid-Mediated Colistin Resistance Gene *mcr-3* in *Escherichia coli*. mBio 2017; 8: e00543–17.

Zankari E, Hasman H, Kaas RS, Seyfarth AM, Agersø Y, Lund O, Larsen MV, Aarestrup FM. J Antimicrob Chemother 2013; 68: 771–7.

Zhang H, Srinivas S, Xu Y, Wei W, Feng Y. Genetic and Biochemical Mechanisms for Bacterial Lipid A Modifiers Associated with Polymyxin Resistance. Trends Biochem Sci 2019; 44: 973–88.

Zhao F, Zong Z. Kluyvera ascorbata Strain from Hospital Sewage Carrying the *mcr-1* Colistin Resistance Gene. Antimicrob Agents Chemother. 2016; 60: 7498–501.

